# Replication of Host *St6galnac1*-Microbiome Association Across Distinct Experimental Conditions in Heterogeneous Stock Rats

**DOI:** 10.64898/2026.10.09.758003

**Authors:** Mohammad Sadegi, Benjamin B Johnson, Thiago Missfeldt Sanches, Gracie Ang, Jennifer Cao, Madison Ambre, Molly Brennan, Olivier George, Rob Knight, Amelie Baud, Abraham A. Palmer, Oksana Polesskaya

**Affiliations:** Department of Psychiatry, University of California San Diego, La Jolla, CA, USA; Department of Pediatrics, University of California San Diego, La Jolla, CA, USA; Department of Computer Science & Engineering, University of California San Diego, La Jolla, CA, USA; Shu Chien-Gene Lay Department of Bioengineering, University of California San Diego, La Jolla, CA, USA; Halıcıoğlu Data Science Institute, University of California San Diego, La Jolla, CA, USA; Center for Microbiome Innovation, La Jolla, CA, San Diego, USA; Centre for Genomic Regulation, Barcelona Institute of Science and Technology, Barcelona, Spain; Universitat Pompeu Fabra, Barcelona, Spain; Institute for Genomic Medicine, University of California San Diego, La Jolla, CA, USA

**Keywords:** sialyltransferase, heterogeneous stock rats, *Paraprevotella*

## Abstract

The gut microbiome is shaped by both environmental and host genetic factors. Identifying specific host loci is essential for understanding the heritability of microbiome-associated traits. The gene *St6galnac1* encodes a sialyltransferase involved in mucin O-glycosylation. Tonnelé et al. (2025) identified significant associations between a locus containing *St6galnac1* gene, and the abundance of genus *Paraprevotella* using cecal samples collected from fasted animals. This association was replicated across three cohorts, and paralleled with association found in humans between a functionally related sialyltransferase gene *ST6GAL1* and *Paraprevotella*.

Given the sensitivity of microbiome analyses to factors including sampling site, feeding status, and analytical framework, independent replication of these findings is important and it is also important to know if the association generalizes across other conditions or is highly specific to certain environments. Here, we performed a replication analysis using stool (not cecal) samples from 174 HS rats that were not fasted prior to collection. Stool samples underwent 16S rRNA gene sequencing, and microbial community structure was analyzed using compositionally aware methods. The rats were genotyped at the top SNP that was associated with *Paraprevotella* in the Tonnelé et al. (2025) study. The association between genotype and *Paraprevotella* abundance was tested using a linear mixed model accounting for sex, cohort, and genetic relatedness. We successfully replicated (p=0.015) *St6galnac1 - Paraprevotella* association.

These findings support the robustness of the *St6galnac1 - Paraprevotella* association across different sampling strategies and experimental conditions.

## INTRODUCTION

The gut microbiome is a complex ecosystem that plays a critical role in host metabolism [1,2,3], immunity [4,5,6], and overall health [7]. While environmental variables such as diet [8] and medication [9] influence the gut microbiome significantly, there is growing evidence that host genetics also shapes microbial composition [10,11,12]. Identifying the specific host loci that influence these interactions is essential for understanding the heritability of microbiome-associated traits. However, the reproducibility of these associations across different experimental contexts and populations is an important concern [13].

In this work, we sought to replicate a previously reported association between *St6galnac1* genotype and *Paraprevotella* abundance in an independent cohort of Heterogeneous Stock (HS) rats under distinct experimental conditions.

Outbred HS rats is a powerful model for dissecting genetics of complex traits including host-microbiome interactions [10]. This population was established from eight inbred founder strains and maintained through outbreeding [14]. In humans, host genetic effects on the microbiome are difficult to disentangle from genetically-driven environmental exposures, such as food choices, meal timing, and social partners. In HS rats experiments the controlled environment minimizes such confounding. The genetic architecture of HS rats enables fine-mapping of associated loci to gene-level resolution.

A recent study identified three replicated microbiome-associated loci in HS rats, the most significant of which involved the gene *St6galnac1* [10], which encodes a sialyltransferase responsible for the synthesis of sialyl-Tn antigens during gut mucin O-glycosylation [15].

Because mucins serve as both a physical barrier and a primary carbohydrate source for gut-resident bacteria [16], genetic variations affecting mucin structure are plausible candidates to alter microbiome **(Fig 1A)**. The associations between this locus and the genus *Paraprevotella* replicated in three out of four cohorts examined by Tonnelé et al. (2025). The variants in strong linkage disequilibrium with the lead SNP were located within the 105 kb region that originated from WN/N founder strain. Sequencing data from WN/N founder strain demonstrated a triplication of *St6galnac1* in this haplotype, suggesting that increased gene dosage is the likely functional mechanism underlying this association. The causal role of *St6galnac1* was supported by associations between the same locus and *Akkermansia muciniphila* and *Muribaculum intestinale* in HS rats, consistent with a mouse knockout study [17] demonstrating that *St6galnac1* directly affects the abundance of these mucin-degrading bacteria. Re-analysis of a human dataset by Tonnelé et al. (2025) also found a suggestive association between *ST6GAL1*, a functionally related sialyltransferase gene, and *Paraprevotella* abundance, pointing to potentially shared host-microbiome interactions across species.

**Figure 1.**
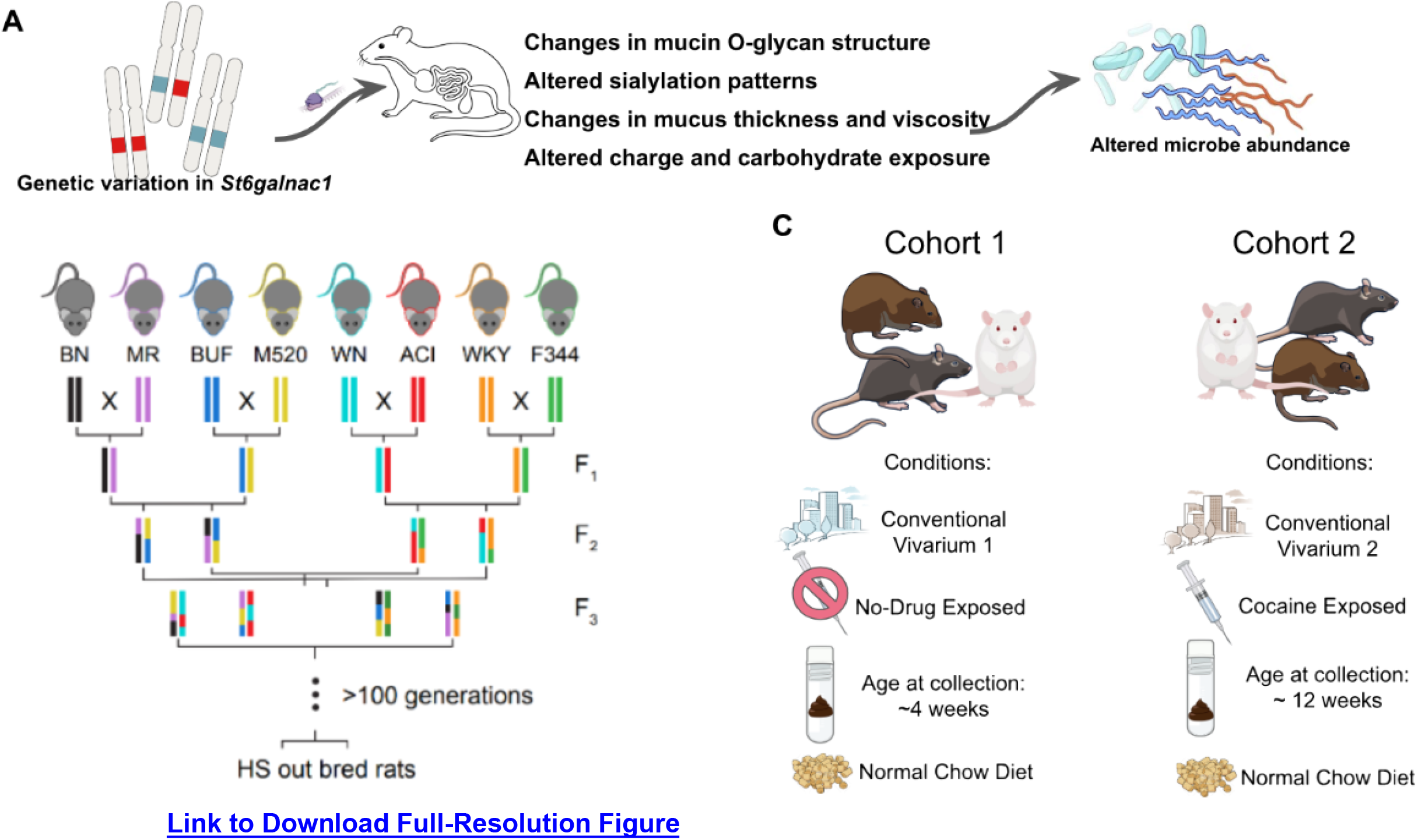
Methodological overview. (A) Study rationale and expectation. (B) Heterogenous stock out bred rat population overview. (C) Overview of the two cohorts used in this study; They differed in age at collection, housing density, vivarium location, and prior experimental history..

To test the robustness of the *St6galnac1*-*Paraprevotella* association in rats, we leveraged two cohorts of HS rats that had been established for independent behavioral studies and derived from the HS West colony of HS rats, which is distinct from the MCW colony used by Tonnelé et al. (2025). We collected samples under conditions distinct from the original study: animals were not fasted prior to collection, stool rather than cecal contents was used, and rats were housed in a different vivarium. We chose to use stool samples and non-fasted animals both because it was more convenient and to better mirror human microbiome research, where using fecal specimens are the standard noninvasive method for analyzing gut microbial communities and are typically collected from participants without controlled fasting [18,19].

To test the reproducibility of this association, we selected rats that have different genotypes for the top SNP reported by [10], measured microbiome composition, and tested the association between genotype at this SNP and *Paraprevotella*. Standard GWAS models assume additive inheritance, but [20] demonstrated that dominance inheritance model accounts for approximately 25% of heritable variance across physiological traits in HS rats and other mammals, and that modeling dominance detects substantially more genetic associations. We therefore performed two analyses: first, we used an additive model, as reported in [10], for replication work. Then we modeled genotype as a categorical variable to allow the mode of inheritance to be determined empirically from the data.

Here we report successful replication of host-genotype at *St6galnac1* locus association with *Paraprevotella* abundance under different experimental and environmental parameters, providing evidence that this host-microbiome signal is robust to substantial variation in sampling strategy and experimental context.

## METHODS

### Animals

All experiments were conducted using outbred N/NIH Heterogeneous Stock (HS) rats (*Rattus norvegicus*) from the colony McwiWfsmAap:HS #155269102, RID:RGD_155269102, generation 101-107. This colony is derived from the colony (NMcwiWFsm, RGD: 13673907) used by Tonnelé et al. (2025); generation 73-80, the rats are separated by approximately 20 generations. The HS rat population was originally established at the NIH in 1984 from eight inbred strains: ACI/N, BN/SsN, BUF/N, F344/N, M520/N, MR/N, WKY/N, and WN/N [14] and the colony has been maintained for over 100 generations to preserve genetic diversity **(Fig 1B)**.

Animals were housed at the University of California San Diego under standard conditions with ad libitum access to water and Teklad 8604 Rodent Diet (Inotiv). Weaning occurred at approximately three weeks of age, at which point pups were separated from their mothers, sex-stratified, and housed in groups of one to four per cage. Each animal was assigned a unique ear punch pattern for identification and implanted subcutaneously with a radio frequency identification (RFID) microchip. Ear punch tissue was collected into individually labeled PCR plate wells and stored for subsequent DNA extraction and genotyping. At this point, animals were assigned to one of two cohorts with distinct housing and experimental conditions. Cohort 1 rats were housed in groups of 2-4 animals per cage and remained in the same vivarium room as the rest of the colony until stool collection at approximately 28 days of age. Cohort 2 rats were transferred to a separate vivarium suite as part of an independent behavioral experiment involving cocaine exposure. During this protocol, animals were housed in pairs or, in some cases, individually. Stool was collected from Cohort 2 rats at approximately 84 days of age, immediately after the conclusion of the behavioral protocol. The differences in housing density, age at collection, and prior cocaine exposure between the two cohorts represent potential sources of variation in microbiome composition and are addressed as confounders in the statistical analyses and discussed in the Discussion section **(Fig 1C)**. All animal procedures were conducted in accordance with the institutional Animal Care and Use Committee (IACUC) of UCSD.

### Rat Genotyping

DNA was extracted from ear punch tissue using the Agencourt DNAdvance kit (Beckman Coulter) via bead-based automated extraction on an Eppendorf epMotion 5075 liquid-handling platform. Extracted samples were normalized to 25.0 ng/μL and randomized prior to library preparation. Libraries were prepared using the 96-plex library preparation kit (TWIST Bioscience). Library quality was assessed via Agilent TapeStation, and sequencing was performed on an Illumina NovaSeq X Plus 25B platform at the UC San Diego Institute for Genomic Medicine (IGM).

Genotypes were called genome-wide using a STITCH-based imputation pipeline, as previously described [21,22], across all rats of the version 8 genotyping round [23]. We selected rats based on the genotype of the top SNP reported by Tonnelé et al. (2025) as the most significantly associated variant with *Paraprevotella* abundance (chr10:101,974,959) because the WN allele has lower frequency (MAF=0.31); genetic selection allowed us to improve power by balancing group sizes across all three genotype classes. From the SNPs called in generation 107 cohort, we identified our additional SNPs in perfect linkage disequilibrium (LD) with this top SNP; only rats with high-confidence genotype calls (INFO score > 0.9) at all five SNPs were retained for further analysis to ensure genotype accuracy. The positions of these SNPs in the mRatBN7.2 (rn7) and GRCr8 assemblies, along with their alleles, are shown in **Table 1**. Animals were classified into three genotype groups based on their consensus genotype across all five SNPs: homozygous reference (BN/BN), heterozygous (BN/WN), and homozygous alternate (WN/WN), where BN refers to the reference allele derived from the BN founder strain and WN refers to the alternate allele derived from the WN/N founder strain, which carries a putative triplication of *St6galnac1*. Each genotype group was defined by concordant genotype calls at all five loci **(Table 1)**.

**Table 1.** SNP positions and alleles.

| Chr:Position<br>(GRCr8) | Chr:Position<br>(mRatBN7.2) | BN/BN<br>(Reference) | BN/WN<br>(Heterozygous) | WN/WN<br>(Alternate) |
| --- | --- | --- | --- | --- |
| 10:102471709 | 10:101972884 | AA | AC | CC |
| 10:102471710 | 10:101972885 | GG | GC | CC |
| 10:102471711 | 10:101972886 | TT | TC | CC |
| 10:102471774 | 10:101972949 | AA | AG | GG |
| 10:102473784 | 10:101974959 | CC | CG | GG |

To maximize statistical power, all available WN/WN rats were included, because this genotype is relatively rare in the HS rats. This pattern is consistent with (Tonnelé et al.), where very few WN/WN animals were observed. Rats with BN/BN and BN/WN genotypes were selected to achieve balanced group sizes. A total of 187 rats were selected for microbiome sequencing across two cohorts **(Table 2)**.

**Table 2.** Microbial sequencing sample size.

|  | BN/BN | BN/WN | WN/WN | Total |
| --- | --- | --- | --- | --- |
| **Cohort 1** | 29 | 33 | 34 | 96 |
| **Cohort 2** | 26 | 58 | 7 | 91 |

### Stool Collection

Stool was collected from rats in Cohort 1 at approximately 28 days of age, and from Cohort 2 at approximately 84 days old. For each animal, identity was verified using an RFID scanner to ensure correspondence between the rat and the pre-labeled collection tube.

Individual rats were removed from their housing cage and placed in a sterile autoclaved cage with a paper sheet covering the bottom. Using sterile forceps, a stool pellet was transferred into a pre-scanned Matrix tube corresponding to that rat’s identification number. Sample tubes were placed on dry ice to preserve sample integrity. Animals were not fasted prior to stool collection. Following collection, each rat was returned to its home cage, and the collection setup was cleaned by replacing the sheet of paper in the cage with a clean sheet for the next rat.

### 16S rRNA Gene Sequencing and Data Processing

Frozen stool samples were transported on dry ice and stored at -80°C until processing. Microbial genomic DNA was extracted using a standard high-throughput bead-beating protocol and normalized to uniform concentration prior to library preparation. Amplicon libraries were prepared using the 16S rRNA Amplification Kit (MiniPCR Bio), a semi-automated adaptation of the Earth Microbiome Project (EMP) standard 16S rRNA V4 workflow, with liquid handling automated across Eppendorf epMotion, Agilent Bravo, Beckman Echo 550, and SPT LabTech Mosquito HV platforms. The V4 region was amplified using EMP primers 515f/806rB in 384-well plates with Platinum II Hot-Start PCR Master Mix (Invitrogen) under standard EMP thermal cycling conditions. PCR products were pooled, purified with the QIAquick PCR Purification Kit (Qiagen), and quantified using the Qubit dsDNA High Sensitivity Assay (Thermo Fisher). Pooled libraries were diluted to 1.0 ± 0.2 nM, spiked with 10% PhiX, and sequenced on an Illumina MiSeq i100 using a 2×151 bp paired-end configuration with custom EMP index primers. Reads were demultiplexed post-run.

QC assessment focused on sequencing depth and diversity structure to evaluate data integrity and identify potential contamination through comparison with blank (negative control) samples. Fecal samples had expected sequence depth of 15,000 to 100,000+ reads, with the majority being above 90,000 reads **(Supplementary Fig. 1A)**. Beta-analysis also displayed separation between samples and blanks; two samples did not meet this requirement and were removed from the study **(Supplementary Fig. 1B)**.

16S rRNA sequencing data are publicly available via Qiita (Study ID: 16160). The downloaded artifact consisted of Deblur 2021.09 processed sequences trimmed to 150 bp, which were imported into a QIIME2 (v2025.10) [23] Python environment for downstream analysis. ASVs were taxonomically annotated against the Greengenes2 2024.09 reference database. Prior to diversity and differential abundance analyses, samples were rarefied to a uniform depth of 60,000 reads; samples failing this threshold were excluded, resulting in a final analytic cohort of 174 animals across two cohorts **(Table 3)**.

**Table 3.** Analytical sample size.

|  | **BN/BN** | **BN/WN** | **WN/WN** | **Total** |
| --- | --- | --- | --- | --- |
| **Cohort 1** | 27 | 30 | 32 | 89 |
| **Cohort 2** | 25 | 53 | 7 | 85 |

ASVs present in fewer than 50% of samples across the full dataset were excluded from analysis, because features with high zero inflation violate the statistical assumptions of linear mixed model, and the data is too sparse to reliably distinguish true biological absence from detection failure, and variance estimates become unstable and untrustworthy.

We confirmed that the ASV in our dataset corresponds to ASV_5163 [10], by a direct sequence comparison. The 150 bp sequence identified in our dataset was identical to ASV_5163 [sequence:TACGGAAGATGCGAGCGTTATCCGGATTTATTGGGTTTAAAGGGAGCGTAGGCGGGAAGTTAAGTCAGCGGTCAAACACGGTTGCTCAACAATCGTTCGCCGTTGAAACTGACTTTCTTGAATGTAGTCAAGGCAGGTGGAATTCGTGGT], ensuring that we are referring to the same organism. ASV_5163 was reported under the Greengenes 2022.10 annotation as *Paraprevotella* by (Tonnelé et al), but more recent annotations under Silva 138.2 represent ASV_5163 as *Prevotellaceae_Ga6A1_group* [d Bacteria;p Bacteroidota;c Bacteroidia;o Bacteroidales;f Prevotellaceae;g Prevotell aceae_Ga6A1_group]. Due to differences in annotation, we refer to this microbe as ASV_5163 in the present manuscript.

### Microbial Diversity Analysis

Within-sample (alpha) diversity was assessed using three complementary metrics: the Shannon diversity index, which accounts for both richness and evenness;the Simpson diversity index, which measures the probability that two randomly drawn reads belong to the same taxon, and observed ASVs, representing raw richness as the count of unique ASV per sample. Group differences in alpha diversity were compared across sex, and cohort.

The 16S rRNA sequencing produces compositional data, where only relative abundances are observed, therefore all beta diversity analyses were performed using compositionally aware methods, which is in keeping with the methods used by Tonnele et al. (2025). Between-sample (beta) diversity was evaluated using Aitchison distance [25], the Euclidean distance applied to centered log-ratio (CLR) transformed abundances. Beta diversity patterns were visualized by Principal Component Analysis (PCA) of the CLR-transformed feature table, and group differences across sex, and cohort were tested using PERMANOVA and PERMDISP on the resulting distance matrix.

### Genotype-Microbiome Association Analysis

Differential abundance of ASVs across genotype groups was assessed using a LMM implemented in the software LIMIX [26]. Genotype, sex, and cohort were included as fixed effects. To account for genetic relatedness between animals while avoiding proximal contamination from the chromosome 10 locus of interest, the genomic relationship matrix (GRM) was constructed from all SNPs, excluding chromosome 10. This approach is called leave-one-chromosome-out (LOCO) [27,28]. Two random effects were modeled simultaneously: LOCO GRM and a cage-level covariance matrix, paralleling the approach by Tonnelé et al. (2025). Variance components were estimated under REML, while model comparison for hypothesis testing (on fixed effects) used maximum likelihood (ML). To estimate the effect of genotype on microbe abundance was we fit three nested models and compared them using a likelihood ratio test (LRT): (1) a categorical model (genotype as a three-level factor) versus null (omitting genotype), to test the overall genotype effect (df = 2); (2) an additive model (genotype as 0/1/2 dosage) versus null, to test the additive trend (df = 1); and (3) the categorical versus additive model, to test dominance effect (df = 1). For visualization, the adjusted phenotype values were computed by regressing out fixed effect contributions from the null model and re-centering to the sample mean for visualization. Because our focus was on replication, and because our replication cohort was too small to provide power for genome-wide discovery we only examined the top SNP identified by [10] as associated with ASV_5163 abundance.

Model used:

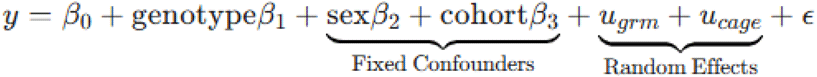

- *y*: Vector of microbe abundance levels.
- β _0_: Intercept.
- genotypeβ _1_: The primary fixed effect being tested.
- sex, cohort: Fixed effect matrices acting as confounders.
- 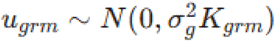: Random effect for host genetics (DGE).
- 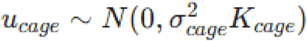: Random effect for the shared cage environment.
- 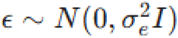: Residual error or “white noise.”

To determine the mode of inheritance at *St6galnac1* locus, a dominance ratio was calculated following the framework described by Cui et al. (2023). We utilized the adjusted mean abundances (residuals from the linear mixed model) for the three genotype classes.

The additive effect (*a*) and dominance effect (*d*) were calculated as follows:

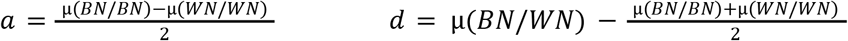

The degree of dominance was then quantified by the ratio *d/a*. Based on this ratio, genetic effects were classified into discrete categories: over-dominant, dominant, or additive.

### Code and data availability

All codes used is available on the Palmer Lab Git repository: GitHub Raw genotypes are available from the UC San Diego Library Digital Collections (Palmer, Abraham A. (2026)). Heterogeneous Stock (HS) Rat Genotypes, Version 8 [23]. In Genotype Data from: NIDA Center for GWAS in Outbred Rats. UC San Diego Library Digital Collections. 16S rRNA sequencing data and associated data and metadata are available in Qiita (Study ID: 16160).

## RESULTS

### Cohort Differences in Microbial Community Structure

As a first step toward testing the reproducibility of the *St6galnac1*-Paraprevotella association, we characterized cohort-level differences in microbial composition, because two cohorts differed in age, housing conditions, and experimental history. To assess whether cohorts should be included as a covariate in downstream analyses, we compared alpha and beta diversity between the two cohorts. Alpha diversity did not differ significantly between groups across in Shannon diversity (p=0.31) and observed ASVs (p = 0.92), but was different under the Simpson test (p=0.021), indicating richness is unchanged, but the evenness/dominance structure differs between cohorts **(Supp Fig. 2A)**. Beta diversity analysis revealed clear cohort separation by PCA, with PC1 explaining 9.1% and PC2 explaining 3.5% of variance **(Supp Fig. 2B)**. PERMANOVA analysis confirmed that this separation was statistically significant (pseudo-F = 12.39, R^2^ = 0.067, p = 0.001), with cohort accounting for approximately 6.7% of total compositional variance. Additionally ASV-level compositional profiles differed between cohorts **(Supp Fig. 2C)**. These results indicate that the two cohorts are compositionally distinct, justifying the inclusion of cohorts as a fixed effect in all subsequent models.

### Sex and Cage Effects on Microbial Community Structure

As a second step, we evaluated contributions of sex and cage to microbial community composition. Both cohorts contained male and female animals (Cohort 1: F=40, M=49; Cohort 2: F=52, M=33), and differed in housing structure in terms of number of cages and animals per cage **(Supp Fig. 3A)**, motivating evaluation of sex and cage as potential confounders.

Beta diversity stratified by sex revealed significant compositional differences in both cohorts (Cohort 1: pseudo-F = 1.91, p = 0.001; Cohort 2: pseudo-F = 2.46, p = 0.001; **Supp Fig. 3B, 3D)**. In Cohort 1, PERMDISP was non-significant (F = 0.50, p = 0.497), indicating that sex differences reflect shifts in community centroids rather than dispersion **(Supp Fig. 3C)**. In Cohort 2, PERMDISP was significant (F = 10.14, p = 0.007), suggesting sex-associated differences reflect both compositional shifts and increased within-group variability **(Supp Fig. 3E)**.

Cage effects were similarly significant in both cohorts (Cohort 1: pseudo-F = 2.72, p = across 19 cages; Cohort 2: pseudo-F = 1.11, p = 0.001 across 62 cages; **Supp Fig. 3F, 3G**). The weaker effect size in Cohort 2 likely reflects variance distributed across a larger number of cages rather than the absence of a cage effect. PERMDISP was not run for cage effects because it would give unreliable estimates of within-group dispersion due to the large number of cages with few animals per cage.

Together, these findings confirm that sex and cage contribute meaningfully to microbial community variation in these cohorts and therefore support inclusion of sex and cage as covariates in all downstream genotype-associated analyses.

### Replication of *St6galnac1*-Microbiome Association

To test for replication of the *St6galnac1* locus-*Paraprevotella* association, we directly tested the effect of genotype on *Paraprevotella* ASV_5163 abundance using a likelihood ratio test (LRT) with full covariate adjustment for sex, cohort, cage and relatedness, we applied a LRT comparing models with and without genotype as a fixed effect. We modeled genotype as an additive dosage (0/1/2), matching the approach used by [10]. This yielded a significant association (χ^2^ = 5.96, df = 1, p = 0.015), confirming the results from [10], namely that this SNP explains a meaningful proportion of variance in *Paraprevotella* abundance after accounting for sex, cohort, cage, and relatedness. Adjusted genotype means revealed that BN/BN animals had substantially higher *Paraprevotella* abundance (2999.1) relative to both BN/WN (1937.7) and WN/WN (1980.6) animals **(Fig. 2A)**. This directional pattern is consistent with the original study [10]. The absolute read counts are higher in the present data, reflecting deeper sequencing, but the relative ordering and significance of the association are replicated. When considering the cohorts separately, the same trend is evident in mean difference between the three genotypes **(Fig. 2B)**.

**Figure 2.**
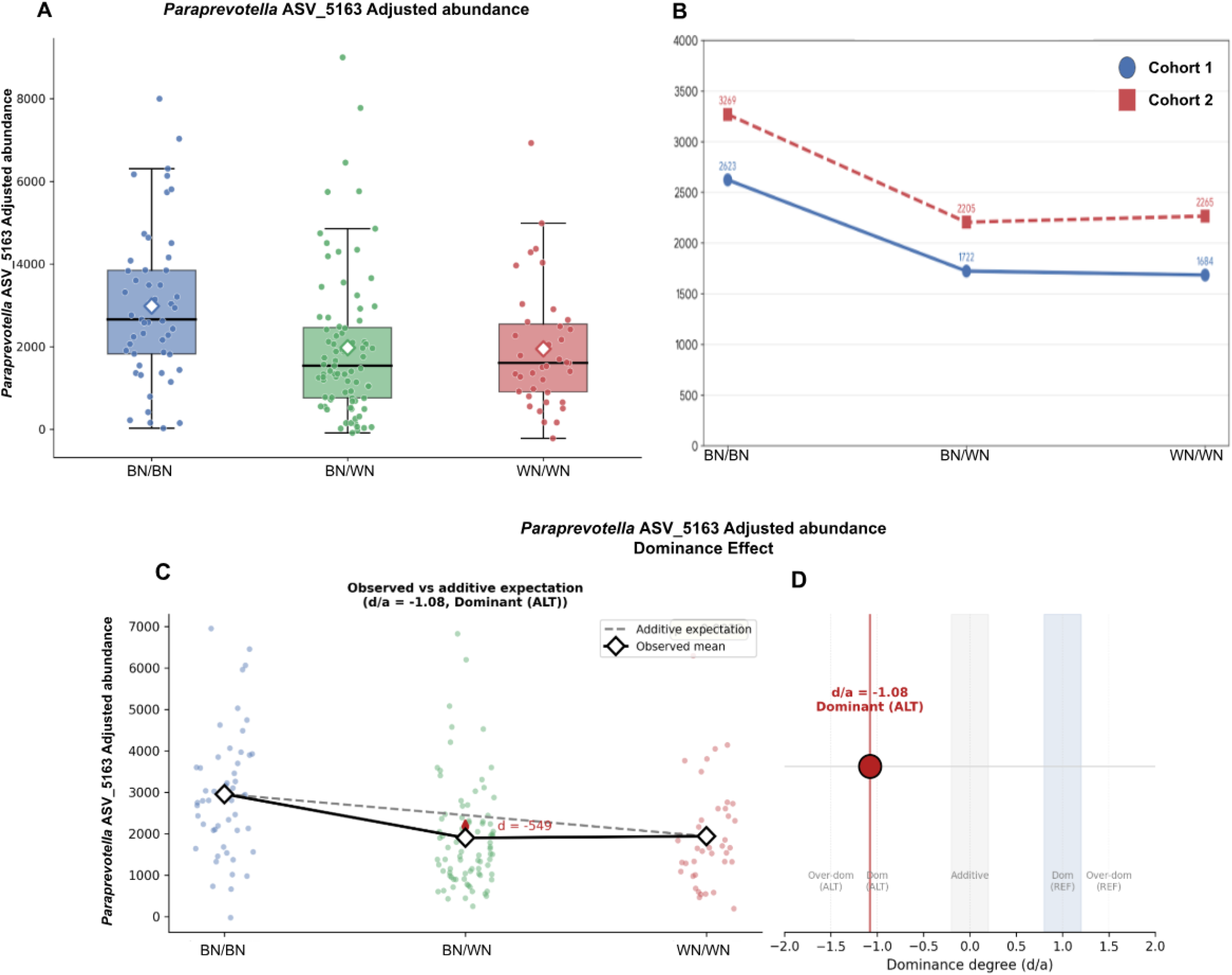
Differential *Paraprevotella* ASV_5163 abundance by *St6galnac1* genotype. (A) Adjusted microbe abundance after accounting for sex, cohort, cage, and genomic relatedness using a linear mixed model. Diamonds indicate adjusted means. (B) Adjusted mean of Paraprevotella abundance by genotype, shown separately for two cohorts. (C) Observed genotype means versus additive expectation plot, showing dominance deviation (d) from the additive midpoint expectation. (D)Cui et al. (2023) Dominance classification; observed value shown in red.

To determine the mode of inheritance at this locus, we additionally modeled genotype as a categorical variable rather than an additive dosage. This allowed the mode of inheritance to be determined empirically from the data. This analysis also found significant association [χ^2^ = 9.27, df = 2, p = 0.0097]. Notably, the near-identical adjusted means of heterozygous and alternate animals relative to the elevated reference group suggested a possible dominance effect of the alternate allele, which we investigated further.

### Dominance Effect of the Alternate Allele at *St6galnac1* locus

The adjusted genotype means suggested a non-additive inheritance pattern, which was not previously reported, with BN/WN animals (1937.7 adjusted abundance) exhibiting ASV_5163 abundance nearly identical to WN/WN animals (1980.6 adjusted abundance) **(Fig 2A)**, rather than falling at the midpoint between the two homozygous groups as would be expected under additive inheritance model. To formally characterize this pattern, compared the categorical genotype model to the additive model by LRT. The dominance term did not reach significance (χ^2^ = 3.317, df = 1, p = 0.069), but suggested the dominance effect. To calculate the magnitude of this effect, we calculated the dominance ratio *d/a*, where *d* is the deviation of the heterozygote from the additive expectation and *a* is the additive effect. A *d/a* of -1 indicates that the heterozygote phenotype is indistinguishable from one of the homozygotes. The observed *d/a* = ™1.08 (*a*= 508.71, *d* = -548.65) indicates complete dominance of the alternate (WN) allele, as the heterozygote phenotype is fully shifted toward the alternate mean rather than the reference mean **(Fig. 2C)**. Using the dominance classification framework that has been previously reported by Cui et al. [20], this pattern falls in the dominant category **(Fig. 2D)**. This pattern was not reported by Tonnelé et al. (2025), whose additive GWAS framework was not designed to detect non-additive inheritance patterns.

### Replication of Secondary *St6galnac1* Locus Associations

Although our primary focus was *Paraprevotella* ASV_5163 association, we examined whether other microbiome associations reported by (Tonnelé et al) for *St6galnac1* locus were also replicated in our dataset. The top SNP for ASV_5163; chr10:101,974,959 (rn7) or chr10:102,473,784 (GRCr8) was also previously associated with *Akkermansia* (ASV_2821) [10], we observed a nominal one-tailed significant association (p = 0.051) in the same direction as in Tonnelé et al. (2025) (**Supp Fig. 4**) using the same sample size shown in **Table 3**, suggesting replication of this association. *Muribaculum* ASV_17008 was also reported by Tonnelé et al. (2025) as associated with *St6galnac1* locus (chr10:101,975,949 (rn7) or chr10:102,474,774 (GRCr8)), but we did not see a significant association (p=0.42 one-tailed).

## DISCUSSION

This study successfully replicated the association between *St6galnac1* locus and *Paraprevotella* ASV_5163 abundance in HS rats, extending the original findings to a distinct experimental context. Critically, this was not intended as an exact methodological replication. The two cohorts that we used differed in age at collection (28 vs 84 days) and prior cocaine exposure. By using stool rather than cecal samples and non-fasted animals, we deliberately tested whether the association persists across biologically meaningful sampling differences, conditions more representative of non-invasive, longitudinal study designs. The fact that the association replicated under these conditions strengthens confidence that the *St6galnac1-Paraprevotella* relationship reflects a genuine host genetic effect rather than an artifact of specific sampling protocols. Furthermore, our rats came from a different HS colony (HS West, generations 101 - 107) than those used by Tonnelé et al. (MCW colony, generations 73-80), providing an additional test of the generalizability of Tonnelé’s findings. We additionally found suggestive evidence of replication of the association between the same locus and *Akkermansia* ASV_2821, consistent with the broader influence of this locus on gut microbial composition. Finally, by modeling genotype as a categorical rather than additive variable, we identified a suggestive previously unreported dominance pattern, in which BN/WN animals phenotypically resemble WN/WN animals rather than showing an intermediate phenotype, suggesting that a single copy of the WN haplotype is sufficient to reduce *Paraprevotella* abundance to levels seen in WN/WN homozygotes. Although the effect did not reach significance (*p* = 0.069), it is notable, because it cannot be detected by the additive models used for GWAS, while known to be common in mammalian genetics [20].

Several limitations need consideration. The two cohorts differed in genotype group distributions, particularly the underrepresentation of WN/WN animals in Cohort 2, which reduced power for certain comparisons. The use of 16S rRNA gene sequencing limits taxonomic resolution to the genus level in most cases, and functional interpretation of the *Paraprevotella* association remains speculative without metagenomic or metabolomic data. Additionally, while cage was modeled as a random effect, unmeasured environmental variables that co-vary with cage or cohort cannot be fully excluded. Finally, the biological mechanism linking *St6galnac1*-mediated mucin O-glycosylation to *Paraprevotella* colonization remains uncharacterized and requires direct experimental investigation.

This study points to several directions for understanding host genetic effects on gut microbiome. To resolve the biological mechanism underlying the *St6galnac1-Paraprevotella* association we will require gut-specific expression data for *St6galnac1*, that would allow us to test whether the WN haplotype influences *St6galnac1* expression in the intestine. The analytical approach that, combining compositionally aware microbiome methods with mixed model quantitative genetics in HS rats can be applied to disease models to reveal how host genetic variation at loci influencing mucosal biology shapes microbiome composition in the context of inflammatory bowel disease, metabolic syndrome, or other conditions where the gut microbiome plays an important role. Identifying genetic variants that simultaneously influence gene expression, glycosylation, and microbial community structure may ultimately help resolve the directionality of microbiome-disease associations that are otherwise difficult to disentangle in observational human studies.

## CONCLUSION

We replicated the association between a locus containing *St6galnac1* gene and *Paraprevotella* ASV_5163 abundance in an independent cohort of HS rats from a distinct colony, using fecal samples from non-fasted animals, confirming that this host genetic effect is robust to differences in sampling site and feeding status. A suggestive association with *Akkermansia* ASV_2821 was also replicated. These results support *St6galnac1* as a reproducible host genetic determinant of gut microbial composition that is robust to substantial variation in experimental conditions, and demonstrate the value of HS rat model for dissecting the genetic architecture of the microbiome.

## Supporting information

All Supplemental Figures

## Acknowledgements

This work was supported by P30DA060810 to AAP. This publication includes data generated at the UC San Diego IGM Genomics Center utilizing an Illumina NovaSeq 6000 and NovaSeq X Plus that were purchased with funding from a National Institutes of Health SIG grant (S10OD026929).

## Author’s contributions

Conceptualization: MS, AAP, OP, AB; Methodology: MS, OP, BJ, TS, GA.; Formal Analysis: MS; Investigation: MS,OP, BJ, TS, GA, JC, MA, MB, OG, RK; Writing (Original Draft): MS; Writing-Reviewing & Editing: MS, OP, AAP, AB; Visualization: MS; Supervision: OP, AAP; Project Administration: OP, RK, AAP; Funding Acquisition: AAP

