## Supplementary material for "Replication of Host *St6galnac1*-Microbiome Association Across Distinct Experimental Conditions in Heterogeneous Stock Rats": All Supplemental Figures

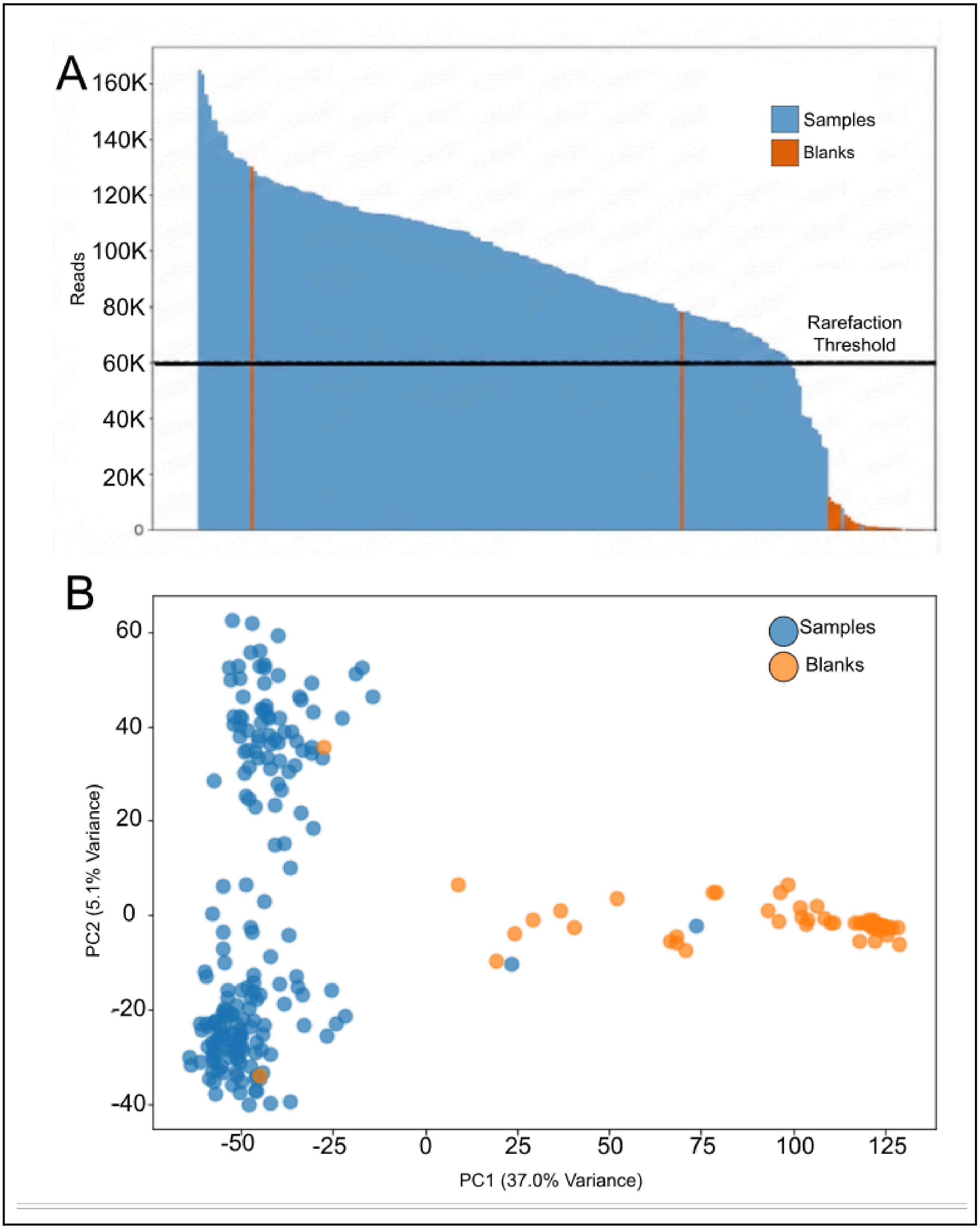
A
160K
140K
120K
100K
80K
60K
40K
20K
0
Reads
Samples
Blanks
Rarefaction Threshold
B
60
40
20
0
-20
-40
PC2 (5.1% Variance)
-50
-25
0
25
50
75
100
125
PC1 (37.0% Variance)
Samples
Blanks

SUPPLEMENTARY MATERIALS

**Supplementary Figure 1. Sequencing quality control metrics.** (A) Sequencing depth across all samples, shown on a log scale. The dashed line indicates the rarefaction depth threshold [60K]; samples below this threshold were excluded from downstream analyses. (B) Aitchison beta diversity comparing fecal samples (blue) and blank negative controls (orange); two samples clustering with blanks were removed from further analysis.

Link to Download Full-Resolution Figure

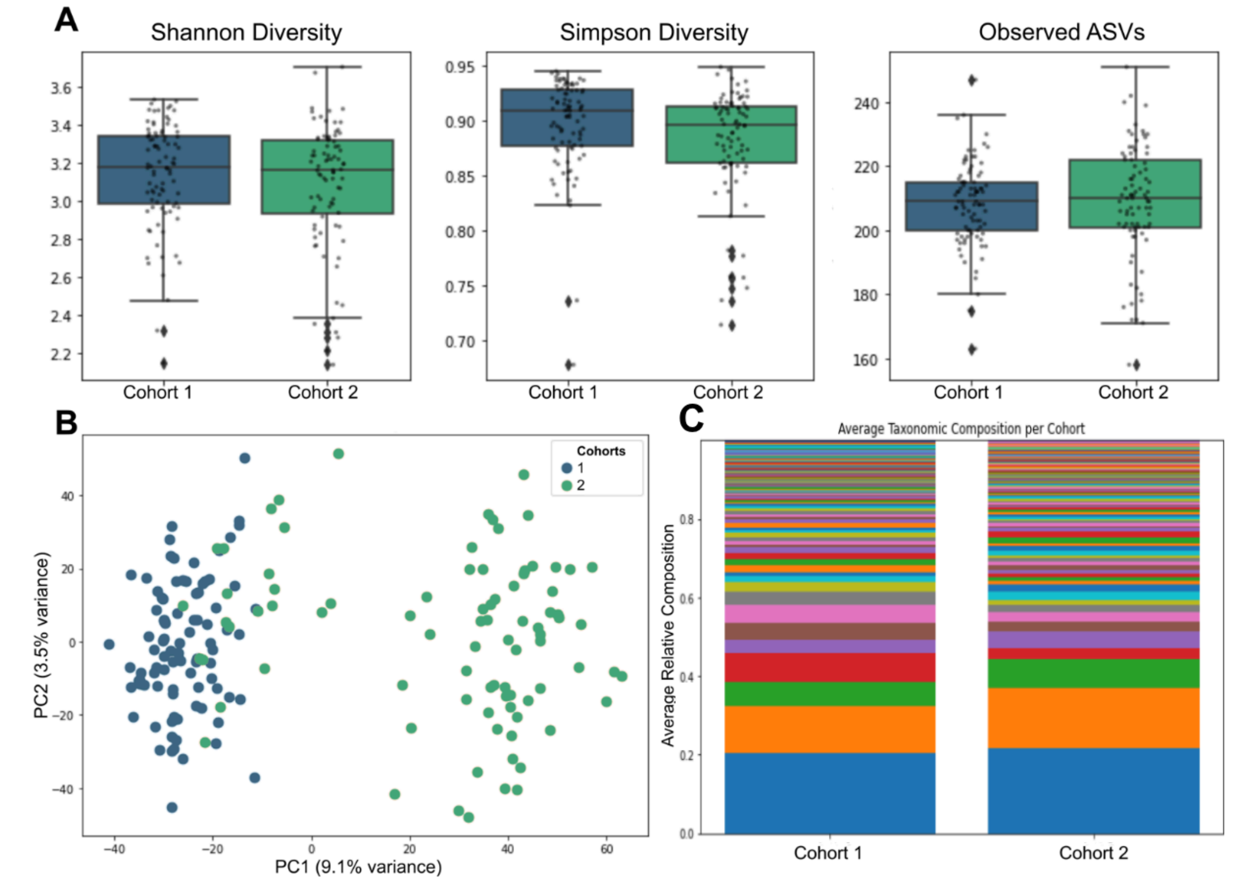


**Supplementary Figure 2. Microbial diversity comparison between cohorts.** (A) Within-sample (alpha) diversity metrics: Shannon diversity index, Simpson diversity index, and Observed ASVs compared between cohorts 1 and 2. (B) Between-sample (beta) diversity assessed by PCA of Aitchison distances. (C) Average taxonomic composition of cohorts; each color represents a unique ASV.

**[Link to Download Full-Resolution Figure]**

A

| | Cohort 1 | Cohort 2 |
|---|---|---|
| Sex | F= 40<br>M=49 | F=52<br>M=33 |
| # of Cages | 19 | 62 |
| Avg Rat per Cage | 4 | 1 - 2 |

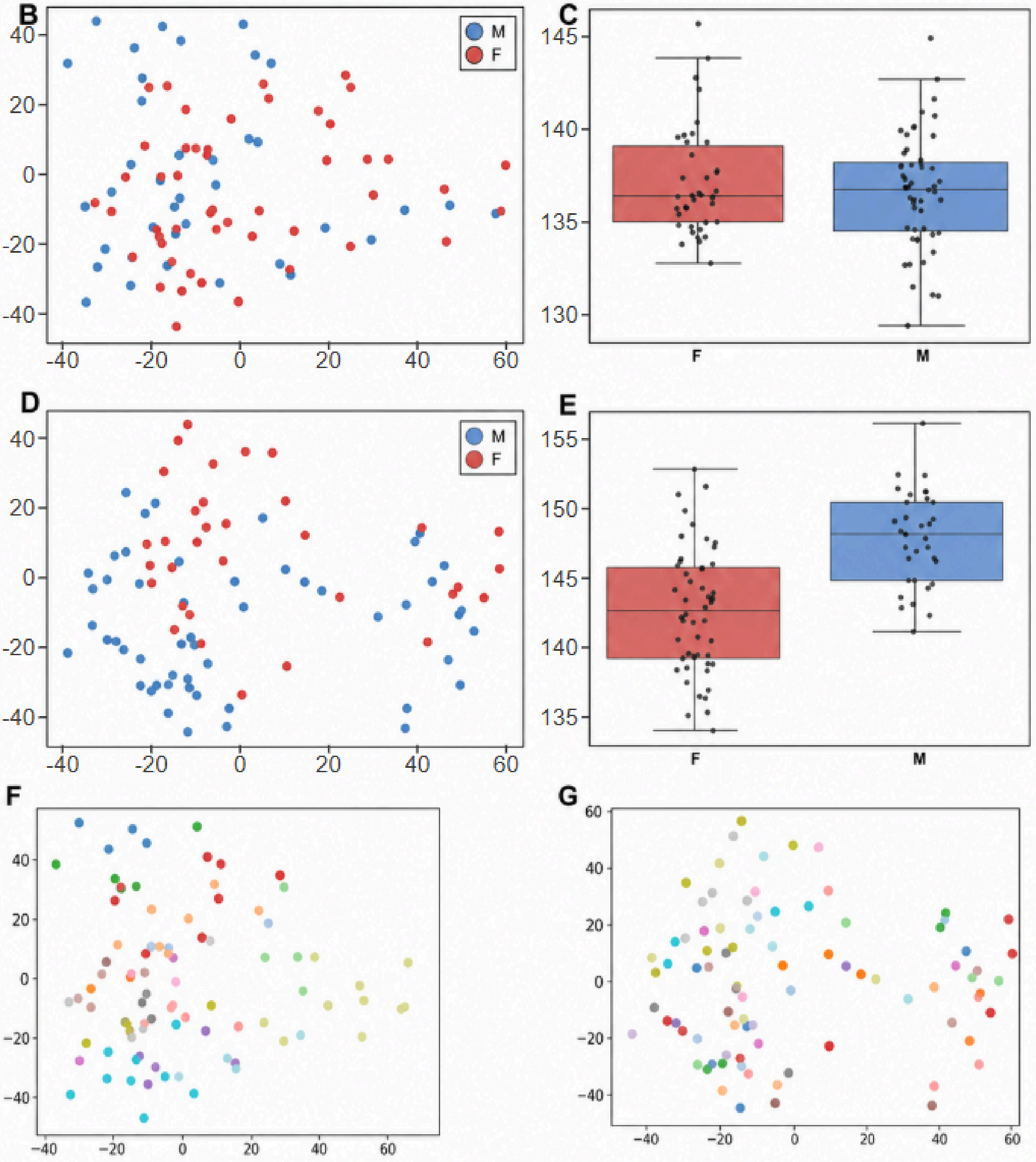


**Supplementary Figure 3. Microbial diversity comparison between sexes and cages** (A)

SUPPLEMENTARY MATERIALS

Sample dimensions. (B) Cohort 1 sex beta diversity. (C)Cohort 1 sex PERMDISP. (D) Cohort 2 sex beta diversity. (E) Cohort 2 sex permdisp. (F) Cohort 1 cage beta diversity (Each color is a cage). (G) Cohort 2 cage beta diversity (Each color is a cage).

**Link to Download Full-Resolution Figure**

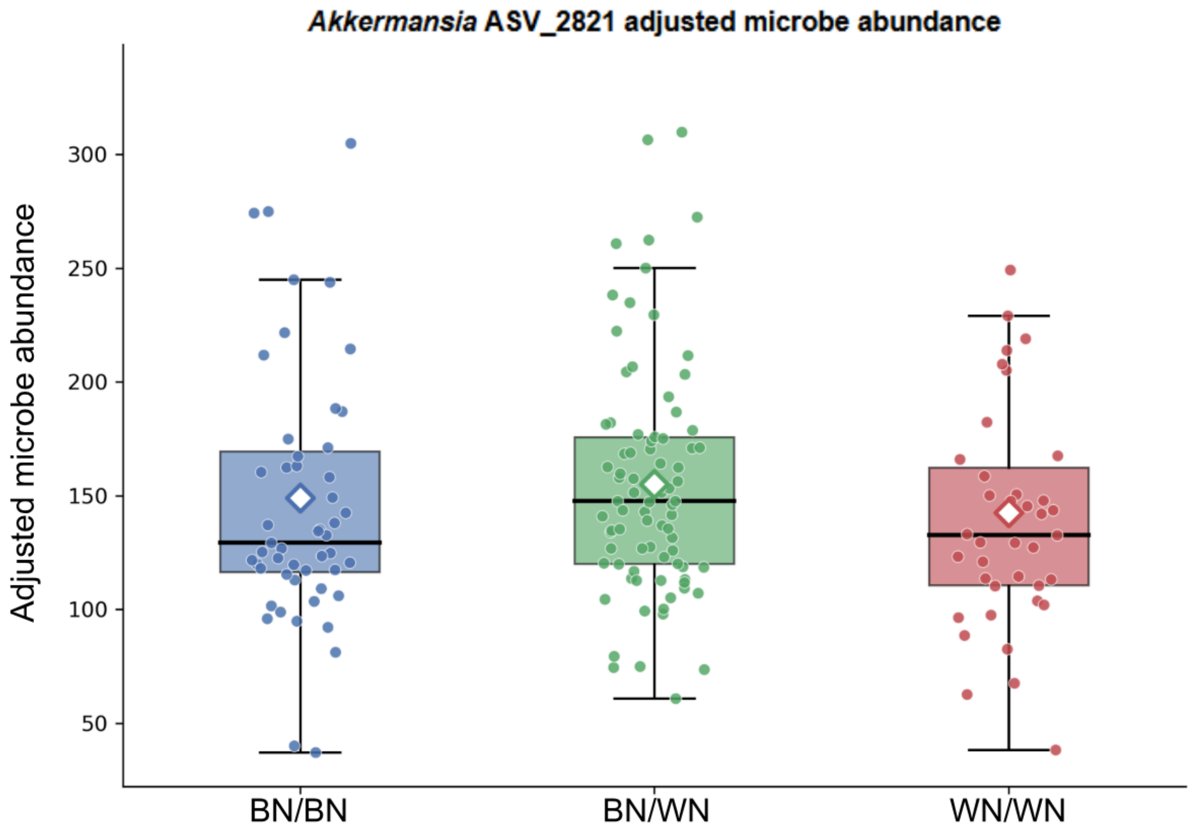


**Supplementary Figure 4. Akkermansia ASV_2821 Adjusted microbe abundance.** Adjusted microbe abundance after accounting for sex, cohort, cage, and genomic relatedness using a linear mixed model . Diamonds indicate adjusted means.

[Link to Download Full-Resolution Figure]
